# Towards Precision Functional Brain Network Mapping in Parkinson’s Disease

**DOI:** 10.1101/2025.07.01.662626

**Authors:** Jake Chernicky, Ally Dworetsky, Sarah Grossen, Emma Carr, Abdulmunaim Eid, Meghan C. Campbell, Caterina Gratton

## Abstract

**Background:** Parkinson’s disease (PD) is a complex neurodegenerative condition that leads to widespread disruption of large-scale brain networks and is further complicated by substantial individual variability in symptomology, progression rates, and treatment response. Consequently, the investigation of individual differences in networks measured via resting state functional connectivity (RSFC) may provide insight. However, most RSFC studies are unable to identify interindividual differences due to poor reliability and group average network definitions. “Precision” RSFC addresses these shortcomings through extended data collection, strict denoising, and individual network definition, but remains untested in PD.

**Objectives:** To evaluate the feasibility and reliability of precision RSFC studies in PD.

**Methods:** We collected >100 minutes of RSFC data from 20 PD and 6 healthy controls participants. We evaluated the level of motion, reliability and stability of RSFC measures in each participant and contrasted these measures between the PD and HC groups, as well as compared to a conventional 5 minutes of RSFC data. In addition, we created individualized brain network measures in PD participants to establish feasibility in this population.

**Results:** Using precision methods, the PD group produced reliable and stable RSFC measures of brain networks of similar quality to healthy controls and substantially better than conventional methods. Individualized network maps from individuals with PD demonstrate differences from group averages and from each other, including in key motor systems.

**Conclusion:** Precision RSFC is feasible and reliable in individuals with PD. This approach holds promise for advancing personalized diagnostics and identifying brain-based biomarkers underlying clinical variability in PD.

## Introduction

Parkinson’s disease (PD) is a multifaceted neurodegenerative condition characterized by a diverse array of pathophysiological changes and symptomatology that transcend its designation as a movement disorder. PD related neuropathology includes progressive degeneration of dopaminergic neurons, coupled with Lewy-body deposits, neurotransmitter deficits, neuroinflammation, and cortical atrophy [1, 2]. However, the precise neurobiological mechanisms underlying the diverse range of specific motor [1, 2], autonomic [3], cognitive [4–6], and affective [7, 8] symptoms remains unclear.

Complicating matters further, people with PD exhibit significant heterogeneity in symptomatology, progression, and treatment response [9–11]. The source of this clinical heterogeneity is widely unexplained [12–15], compromises quality of life, and exacerbates the challenges in identifying efficacious therapies. Thus, successful treatment necessitates individualized evaluation and personalized interventions tailored to each patient [16].

One explanation for the extensive range of symptoms associated with PD is that neuropathology disrupts communication among a diverse array of interconnected brain regions or “networks”. Resting-state functional connectivity (RSFC) is a technique to non-invasively evaluate large-scale brain connectivity [17] and networks [18] and has demonstrated sensitivity to PD [19–21]. In fact, network dysfunction relates to measures of neuropathology (e.g., serum and CSF biomarkers) [22–27] and clinical manifestations including cognitive deficits [28–30] and psychiatric symptoms [31–33].

Thus, previous RSFC studies offer insight into the network characteristics of PD and their relationship to clinical manifestations [15].

However, most RSFC studies use group-level approaches, pooling data from numerous individuals, and thereby reducing clinical relevance and compromising statistical power at the individual level [34]. Because these studies necessitate large numbers of participants [35], data collection is often limited to 5-10 minutes of RSFC data, which is insufficient to reliably identify individual-level brain network maps [36–39]. Other methodological concerns such as in-scanner head motion are also often inadequately addressed, further limiting reproducibility at the individual level [40–42].

“Precision” RSFC, a novel approach centered on the reliable detection of individual differences rather than group-average outcomes, seeks to address these shortcomings through extended data collection (>40 minutes of RSFC collected over multiple sessions), strict denoising for motion and other physiological confounds, and advanced network definition techniques [39, 43, 44]. Prior studies in healthy controls demonstrate that precision RSFC enables reliable and stable [39, 45] individual-level network measures that are ordinarily obscured by conventional methods. These approaches have uncovered meaningful individual-level differences in network topology [46, 47] that correlate with psychopathology [48]. The advantages of precision RSFC may be particularly relevant for studies focused on the subcortex [49, 50], where most MRI has lower signal-to-noise ratio (SNR) [51, 52], but alterations in connectivity may be most evident in PD. Therefore, precision RSFC may be key to uncovering the functional correlates of clinical heterogeneity in PD. With highly reliable individualized network maps, we could systematically document variations in PD from healthy controls, elucidate the mechanisms underlying clinical heterogeneity, capture longitudinal disease progression, and identify biomarkers for the early detection, subtyping, and prognosis of individuals with PD.

However, precision RSFC is untested in individuals with PD and is complicated by methodological concerns such as fatigue during long scanning sessions, increased head motion, and possible inter-session instability due to neuropathology and/or symptomatology. These concerns must be addressed to determine the viability of precision RSFC in PD.

To establish the feasibility and reliability of precision RSFC methods in PD, we collected over 100 minutes of RSFC data from 20 individuals with PD and 6 healthy controls. First, we assessed feasibility by comparing 1) the number of sessions required to complete data collection and 2) in-scanner head motion between groups. To evaluate reliability, we compared test-retest estimates from 40-minutes of data, the minimum for reliable individual-level network measures [37, 39, 53], to those from typical (5-minute) data collection. Data were concatenated across sessions per standard practice [45, 53, 54]. Next, we examined intra- and inter-session similarity of individual correlation matrices, assessing both stability and functional distinction. Finally, we present a proof-of-concept evaluation of individual-level network maps in PD to contrast features identified across two participants.

## Methods

### Participants

Participants were recruited from two ongoing longitudinal studies that incorporate multimodal imaging, biospecimen analysis, and clinical follow-up from PD and healthy control participants [15, 25, 55]. Eligible participants were ≥50 years old, English-speaking, and ≥12 years of education. PD participants met clinical diagnostic criteria; HCs had no neurological disorders and no first-degree relatives with PD. Exclusion criteria included other neurological conditions, schizophrenia, bipolar disorder, dementia (CDR ≥1 [56], MMSE <24, or clinical exam), history of moderate/severe head injury, or MRI contraindications (e.g. deep brain stimulation). Motor severity was assessed in the OFF-medication state using the Unified Parkinson Disease Rating Scale motor subscale (UPDRS-III) [57] with a movement disorder specialist after overnight withdrawal of PD medications. When the movement disorder specialist was not available, participants were video-recorded completing the UPDRS-III and scores were imputed per established methods [58]. The Washington University in St. Louis Institutional Review Board approved all procedures; all participants gave written informed consent.

### MRI Acquisition

Participants completed 3-5 scan sessions over a period ranging from 6 weeks to 7 months (mean = 12.93 weeks, median = 12.07 weeks) after overnight withdrawal of PD medications. MRI scans were acquired with a 3T Siemens Prisma scanner using a 20-channel head coil. MRI sequences included: T1-weighted (TR=2400ms, TI=1000ms, TE=3.18ms, FA=8°, TA=8:09, 0.9mm voxels); T2-weighted (TR=3200ms, TE=294ms, TA=5:38, 0.9mm voxels); echo-planar imaging BOLD (3-10 runs per session; TR=800ms, TE=26.6ms, FA=61°, TA=5:39, 3mm voxels, multiband factor of 4; 416 volumes); and GRE field map images (TR=400ms, TE1=4.92ms, TE2=7.38ms, FA=60°, TA=0:54, 3mm voxels, 36 slices, FoV read=192mm or TR=509ms, TE1=4.92ms, TE2=7.38ms, FA=60°, TA=1:16, 3mm voxels, 48 slices, FoV read=213mm, depending on prior study data collection). Two PD participants’ data contained field map acquisition errors that hindered RSFC processing but were included in feasibility analyses.

### Behavioral QC

During scanning, participants were closely monitored for drowsiness (via eye tracker) and body movement (e.g. tremor). Participants completed the Stanford Sleepiness Scale [59] after each run. Functional runs were excluded if participants had sustained eye closure or movement artifacts for ≥40% of frames or reported high sleepiness (score >5). Two PD participants were excluded from movement and RSFC analyses due to drowsiness but were considered informative to the feasibility of the data collection paradigm. Two of 4 sessions collected from participant P020 were excluded entirely due to sustained tremors.

### MRI Preprocessing

MRI data were organized in BIDS format [60–62] and preprocessed using fMRIPrep v20.2.0 [63]. Structural images underwent intensity correction, skull stripping, and spatial normalization. Functional images were corrected for field distortions and motion, aligned to the anatomical image, and resampled to MNI space. Subsequent processing for RSFC measures followed Power 2014 [64], and included nuisance regression (motion parameters, white matter, CSF, global signal), temporal filtering (0.009–0.08 Hz), and motion censoring using filtered framewise displacement (fFD > 0.1 mm) [64]. Censored volumes were interpolated prior to filtering and removed before connectivity analysis. Further details are available in **Supplemental Methods**.

### Motion Calculation

To quantify head motion, we calculated the amount of data excluded during motion censoring based on filtered framewise displacement (fFD > 0.1 mm). Retained and censored volumes were converted to minutes of scan time for comparison across groups. Group-level differences in total retained and censored scan time were assessed using Welch two-sample t-tests.

### Functional Connectivity

RSFC was computed as Pearson correlations between denoised, motion-censored BOLD time series. ROIs were defined using the Seitzman 300 ROI atlas [65] (**Figure 2A**), which includes subcortical and cerebellar coverage relevant to PD [15, 66]. To account for potential atrophy, ROIs were masked using each participant’s gray matter segmentation from FreeSurfer. All voxels contained within the mask were retained in the cortex and cerebellar ROIs; subcortical ROIs were masked at a more conservative threshold of 50%. ROIs with <25% of their original volume were excluded, resulting in individualized ROI sets (mean ROIs excluded: PD = 1.95, range = 0–7; HC = 1.17, range = 0–3). Final RSFC estimates were represented as 300 × 300 correlation matrices for each participant.

### Test-Retest Reliability

Within-participant test-retest reliability was assessed by randomly shuffling and concatenating BOLD runs to create two independent RSFC estimates per participant. The first 70 minutes of data served as a “high-confidence” reference; remaining data were used to compute RSFC estimates across increasing durations, with correlations calculated at each 1-minute increment. This process was repeated 50 times per participant with different randomly selected segments. Reliability was evaluated within cortex, subcortex, and cerebellum by isolating functional connectivity associated with each structure. Paired t-tests compared reliability between 5-minutes and 40-minutes of data.

### Inter-session Similarity

To examine RSFC consistency across days, we computed pairwise Pearson correlations between functional connectivity matrices from sessions with ≥25 minutes of usable data. Within-participant similarity was defined as the average similarity across session pairs from the same individual; across-participant similarity was calculated between sessions from different individuals. Group-level comparisons were made using Welch-corrected independent-samples t-tests. To evaluate how data quantity influences reliability, we recomputed functional connectivity matrices from the first 5, 10, and 25 minutes of each session, and compared resulting within-participant similarity scores using paired t-tests.

### Individualized Network Mapping

To demonstrate the utility of precision RSFC for mapping individual network organization, we generated seed-based and data-driven functional networks for two PD participants (P001, P026). RSFC time series were projected to grayordinate space (87,129 total; cortical, subcortical, cerebellar) [39, 67, 68] and smoothed (3 mm FWHM). Seed-based connectivity maps were generated using a motor cortex (M1) seed. Data-driven networks were identified using Infomap (v2.2.0) [69], following the individualized parcellation pipeline described by Lynch et al. (2024) [48]. Communities were computed at a graph density of 0.001 and labeled based on spatial and functional similarity to canonical networks. Low-confidence or small parcels (<50 grayordinates) were reviewed and potentially excluded. Two of 60 communities in P001 and 1/61 in P026 were reassigned; 9/60 (P001) and 3/61 (P026) were excluded due to lack of network structure, likely due to poor SNR. The same mapping pipeline was also applied to a group-averaged PD connectivity matrix.

### Data Sharing

The data supporting the findings of this study will be made available upon reasonable request to the corresponding authors. The RSFC processing code is available on GitHub. (https://github.com/GrattonLab/Chernicky_2025_PD_feasibility.git).

## Results

### Participants

PD (N = 20) and HC (N = 6) groups were similar in age, education, and sex (ps > .05); motor severity, as measured by the UPDRS-III, was significantly worse for the PD group, (p<0.05; see **Table 1**).

**Table 1:** Participant Characteristics.

| **Table 1: Participant Characteristics** |  |  |
| --- | --- | --- |
|  | **PD** | **HC** |
| **n** | 20 | 6 |
| **Sex (M/F)** | 8/12 | 2/4 |
| **Age** | 67.4 ± 6.2 | 73.0 ± 7.2 |
| **Education (years)** | 16.2 ± 2.0 | 16.2 ± 1.8 |
| **Baseline CDR-SB (0/0.5)** | 16/4 | 6/0 |
| **Duration PD Motor Sx** | 9.0 ± 5.2 | — |
| **UPDRS-III (OFF Meds Total)** | 25.7 ± 8.9* | 9.1 ± 4.9 |
| **Mean sessions collected** | 4.05 | 4.00 |
| *p<0.05 |  |  |

### Sufficient low-motion RSFC data can be collected from people with PD

Even subtle movement can introduce RSFC bias [40, 70], raising concerns about the feasibility of precision RSFC studies in PD. To address this, we applied conservative motion censoring (fFD < 0.1 [64]); see *Methods*), and calculated the amount of remaining data in each participant.

On average, PD participants retained 108 minutes (91.5% ± 6.9) of data, comparable to HC (113 min, 92.3% ± 9.5%; ps > 0.6; Figure 1). Censored data (PD = 10 min; HC = 9 min; p = 0.79) and session count (PD = 4.05 ± 0.7; HC = 4.00 ± 0; p = 0.19) did not differ (see also **Supplemental Figure 1**). Thus, even with stringent motion control, PD participants provided sufficient data for precision RSFC.

**Figure 1:**
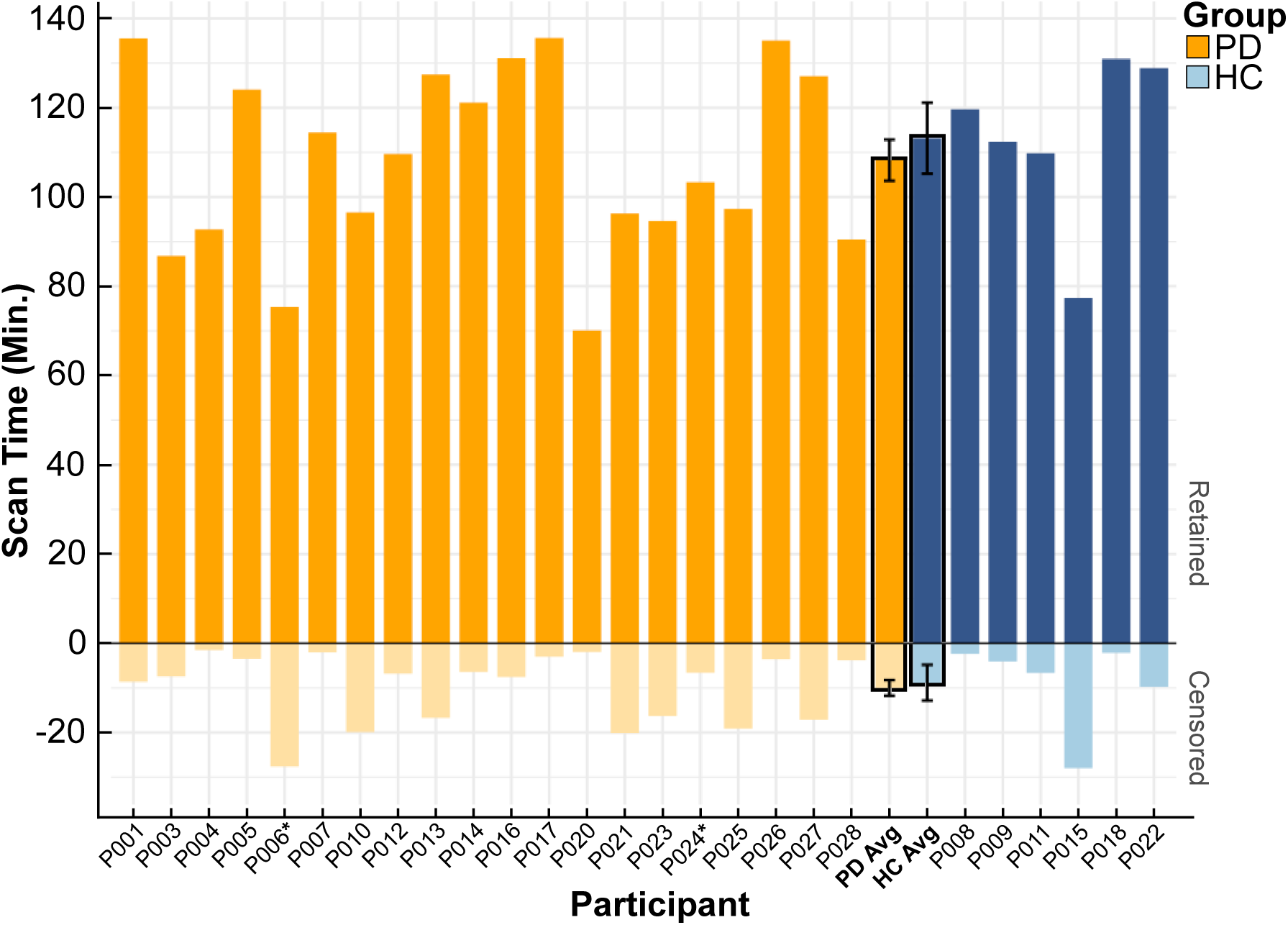
Total Scan Time Retained vs. Lost to Motion Contamination. PD (orange) and healthy control (blue) groups retained and lost similar amounts of data after addressing motion contamination. High motion frames (defined as filtered-framewise displacement of >0.1mm, see Methods) were identified and censored from analyses. *Participants P006 and P024 were excluded from further RSFC analyses due to field-map acquisition errors.

### PD RSFC data achieves high reliability with sufficient data

Next, we examined individual-level RSFC reliability. A sub-sampling analysis (**Figure 2A**) compared two benchmarks: a conventional 5-minutes of data (see Gratton et al., 2018 [15] Supplemental Table 1) and a precision-level 40-minutes of data collection. We report results in the cortex, subcortex, and cerebellum separately (**Figure 2**, see **Supplemental Table 1 & Supplemental Figure 3** for more structures).

**Figure 2:**
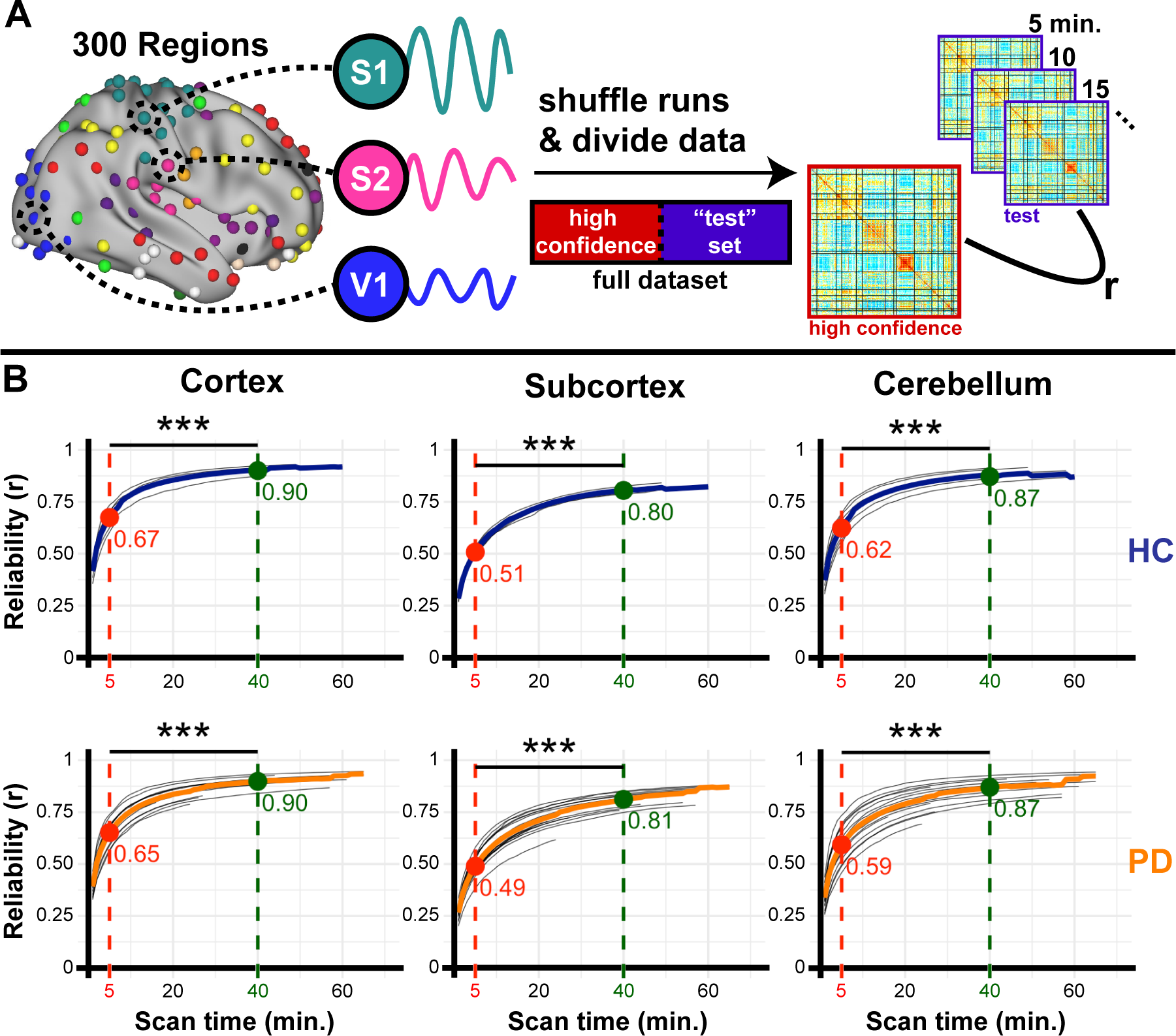
Test-retest reliability improves with scan time. **A)** Run-level fMRI data from ROIs across the brain were randomly shuffled and split into ‘high confidence’ (70 min.) and test (remainder) subsets. Networks calculated from the high-confidence and increasingly-data rich test sets were compared. **B)** Reliability improved with the addition of scan time in ROIs within the cortex, subcortex, and cerebellum at similar rates in both HC (blue) and PD (orange). Each gray line depicts one participant; thick lines indicate the mean. Dashed vertical lines mark 5 min. (red; typical amount of RSFC included in many past studies) and 40 min. (green; minimum amount of RSFC data included in precision fMRI studies).

In the cortex (**Figure 2B, column 1**), PD participants showed moderate reliability with 5 minutes of data (r = 0.65 ± 0.05), which improved substantially with 40 minutes (r = 0.90 ± 0.03), matching the pattern seen in the HC group (5 min: r = 0.66 ± 0.04; 40 min: r = 0.90 ± 0.02). Reliability gains were even larger in the subcortex (PD: r = 0.49 → 0.81, Δ = 0.32; HC: r = 0.51 → 0.80, Δ = 0.29; **Figure 2B, column 2**) and cerebellum (PD: r = 0.59 → 0.87, Δ = 0.28; HC: r = 0.62 → 0.87, Δ = 0.25; **Figure 2B, column 3**). Overall, extended scan time markedly improves the reliability of RSFC estimates across all brain regions, especially those relevant to PD pathology.

### PD RSFC data is stable and distinct across sessions— given sufficient data

Next, we asked whether RSFC network measures remain consistent across sessions. With 25 minutes of data, within-participant similarity of networks was high in PD (r = 0.70 ± 0.09), closely matching healthy controls (r = 0.71 ± 0.07), and aligning with reliability estimates (**Figure 2**). In contrast, across-participant similarity of RSFC was substantially lower (PD: r = 0.32 ± 0.09; HC: r = 0.33 ± 0.06; ps < .0001), indicating distinction in network patterns across individuals. This pattern held across regions, though estimates varied with differences in SNR [50] and reliability (see **Supplemental Figure 2**). These findings demonstrate that individuals with PD exhibit stable and distinct RSFC profiles across sessions, comparable to healthy controls.

Differences in stability estimates across sessions may be driven more by data reliability than by true day-to-day fluctuations in connectivity profiles. To test this, we conducted analyses using sub-sampled data from the first 5, 10, and 25 minutes of each session (**Figure 3B; see also Supplemental Table 2**). In both groups, stability improved with scan duration: it was lowest at 5 minutes (PD: r = 0.38 ± 0.10; HC: r = 0.42 ± 0.09), increased at 10 minutes (PD: r = 0.51 ± 0.11; HC: r = 0.54 ± 0.09), and peaked at 25 minutes (PD: r = 0.70 ± 0.09; HC: r = 0.71 ± 0.09). Paired t-tests confirmed significant gains at each step in both groups (all ps < .005).

**Figure 3:**
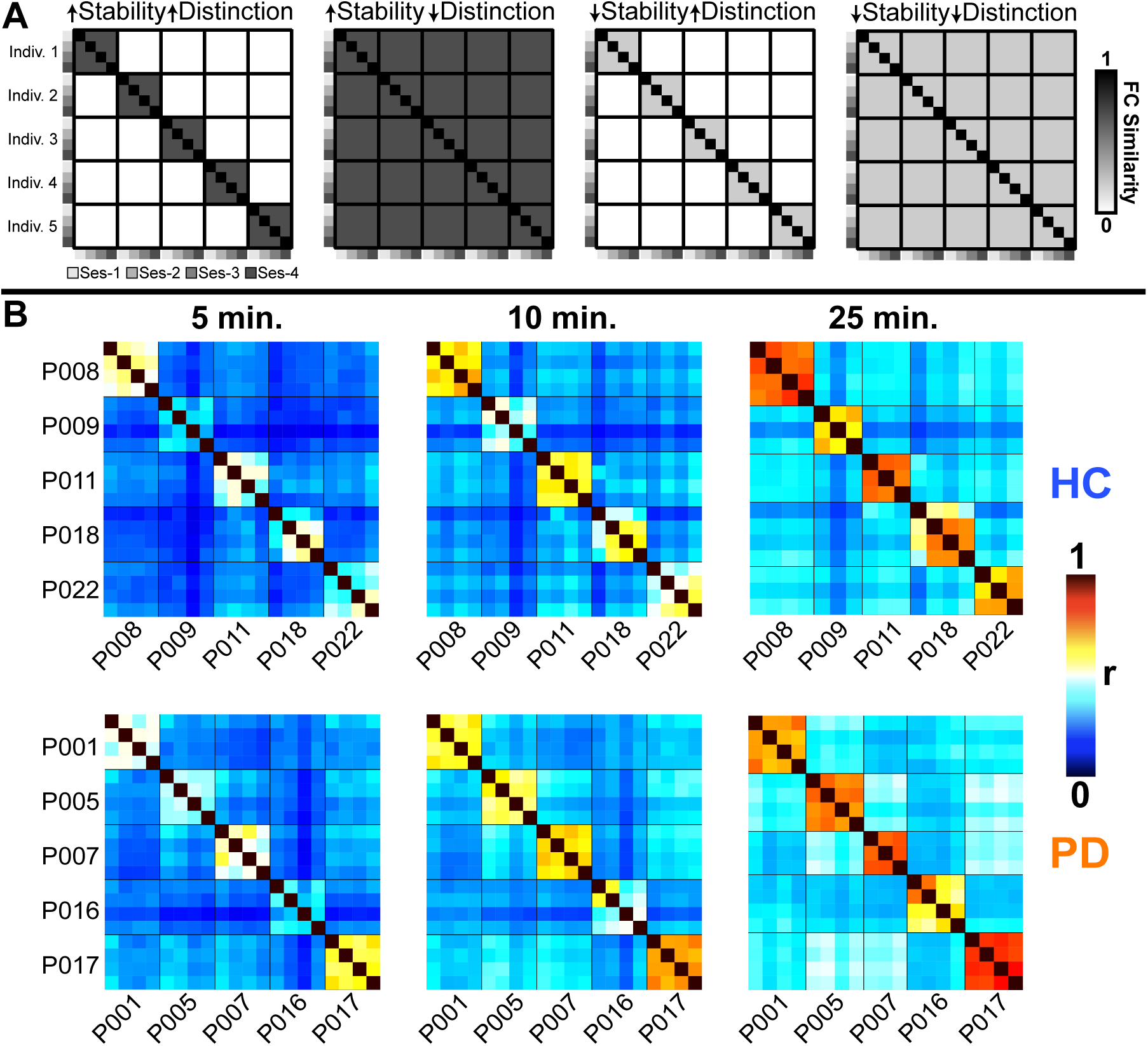
Session to session stability for RSFC. **A)** This figure depicts cartoons with different scenarios for similarity in RSFC across sessions. (Left plot) High similarity in RSFC across sessions from the same person and high distinction across people would indicate RSFC can be used to identify reliable individual differences. (Column 2) High similarity in RSFC across sessions in a person with low distinction across people suggests the presence of a shared and stable network profile. (Column 3) Low similarity in RSFC across sessions with distinction across people is consistent with an unstable but unique network pattern. (Column 4) Low stability in RSFC across sessions within a person and low distinction across people is consistent with unreliable and inconsistent networks (right plot). **B)** Data from ten representative participants in this dataset demonstrate that both PD and healthy control groups exhibit increased session-to-session RSFC stability and distinction as scan time increases, reflecting enhanced measurement reliability with longer data acquisition. A subset of data is shown for clearer visibility, but full group results are consistent and reported in **Supplemental Figure 4**.

### Precision RSFC Enables Individualized Functional Mapping in PD

A key aim of precision RSFC in PD is to identify individual-specific network features that may be linked to neuropathology and clinical manifestations. With reliability and stability established, we present RSFC maps from two PD participants to illustrate how precision fMRI captures personalized connectivity patterns.

**Figure 4A** highlights differences in motor cortex connectivity. In participant P001, the seed location in M1 (white dot) shows broad connectivity with the ventral motor strip consistent with a location in the somatomotor face network [39, 71, 72]. In contrast, in P026 this location showed distinct connectivity with areas resembling the somatomotor cognitive action network (SCAN) [72], indicating that participant-specific differences emerged in motor areas relevant to PD pathophysiology [73].

**Figure 4:**
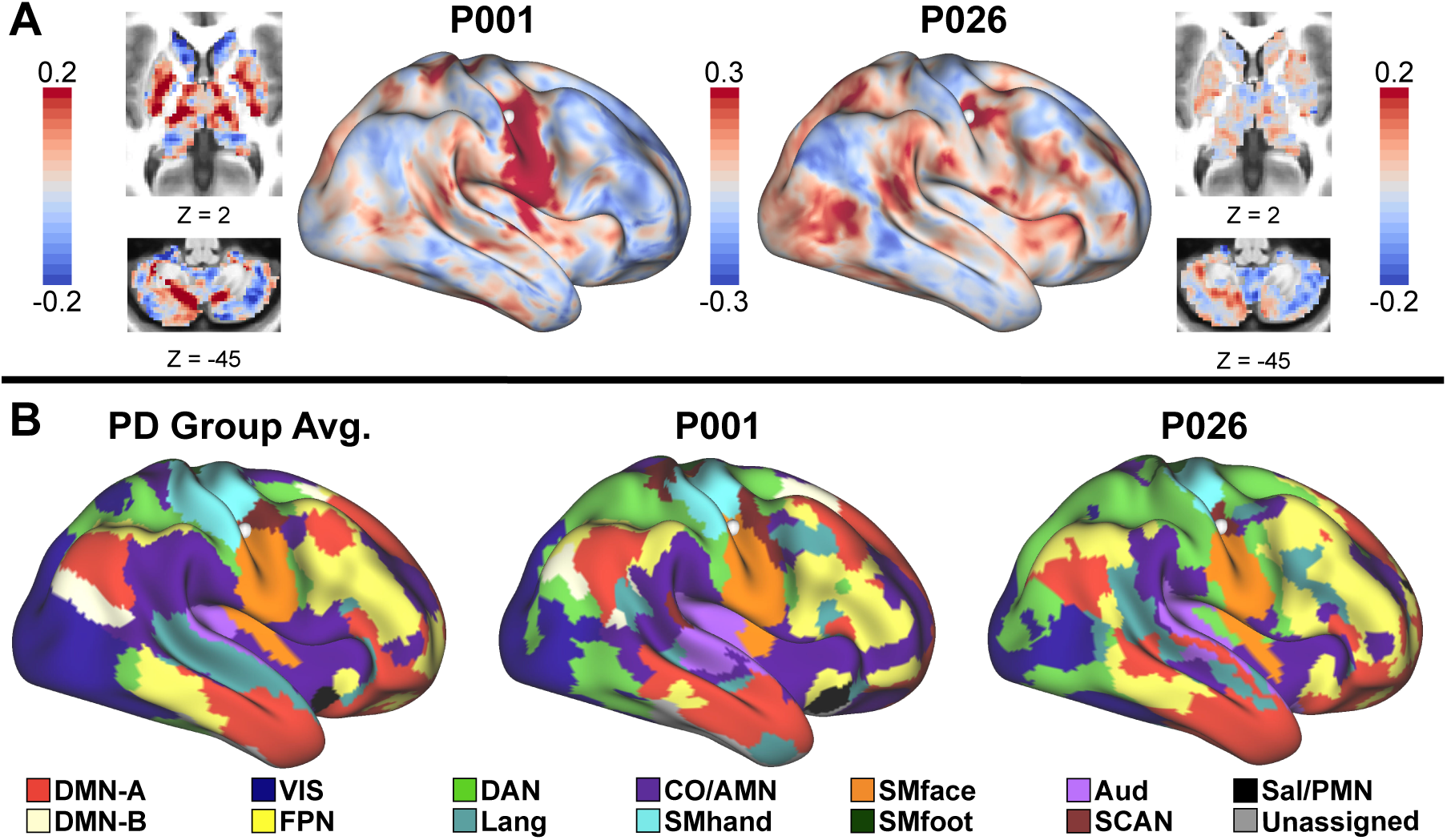
Whole-brain network maps demonstrate individual differences in RSFC. **A)** RSFC of the M1 seed (white dot) differs in P001 and P026. **B)** Data-driven estimation of brain networks in the group (left) and two PD participants (middle and right) based on RSFC (120 min.). Network topology from PD participants differ from each other and the group average (e.g. note differences in lateral prefrontal cortex and near the location of the white seed from panel A).

**Figure 4B** extends this comparison across the cortex, contrasting individualized and group average network parcellations. While both individuals exhibit general similarity to the group in the broad distribution of networks, clear individual-specific deviations in network layout are evident, consistent with past precision RSFC studies [46, 74]. Notably, the lateral prefrontal cortex shows marked differences in the size and position of the cingulo-opercular, language, dorsal attention, frontoparietal, and default mode networks. These individualized patterns underscore the potential of precision RSFC to explore whether network variation relates to clinical heterogeneity in PD.

## Discussion

We examined the feasibility, quality, and reliability of precision RSFC approaches in PD, with the goal of using these techniques to investigate individual differences in large-scale brain systems in PD. PD participants tolerated precision RSFC data collection well and produced data of similar quality and quantity to healthy controls and past precision RSFC studies— with equivalent levels of motion, test-retest reliability, and intersession similarity. Subsequently, we presented evidence that network maps generated from participants with PD show relevant individual variation.

### Precision RSFC is viable in PD

Our findings support the viability of precision RSFC data collection in PD. We were able to acquire extended RSFC data, across multiple sessions, in both groups. Two participants were excluded due to acquisition errors and two due to drowsiness. Two additional sessions from a PD participant were excluded due to sustained tremor throughout the scans. These reflect common challenges of RSFC research in PD rather than specific limitations of precision fMRI [75, 76]. Despite this, the precision RSFC approach of collecting data across multiple scan sessions allowed us to retain the participant with tremor, demonstrating a clear advantage over typical single-session approaches.

Perhaps the most anticipated challenge of precision RSFC in a PD sample is the difficulty of overcoming in-scanner head motion, given that motion is a common and pervasive confound in RSFC studies [42, 70, 77] and inevitable in a PD sample. Despite elevated UPDRS-III scores, all PD participants contributed >40 minutes of low-motion RSFC data. Motion varied between individuals but remained consistent within individuals across sessions, in line with prior work suggesting motion is trait-like [78, 79]. Through denoising strategies and the ability to spread data collection across multiple sessions, precision RSFC was able to overcome limitations from low-data studies and improve participant retention.

### Precision RSFC yields reliable and stable measures of brain networks in PD

Our test-retest results highlight three key findings. First, individuals with PD achieved reliability levels comparable to healthy controls and prior precision RSFC studies [39]. Second, the reliability and stability of RSFC estimates based on >40 minutes of data substantially outperformed conventional 5-minutes of data. Third, precision RSFC offers a distinct advantage in boosting reliability (r = 0.49 at 5 min, 0.81 at 40 min., a 65% increase) in subcortical regions with poor SNR, enabling more reliable assessments of areas implicated in PD pathology. This opens possibilities for early detection, linking connectivity patterns to clinical features, and measuring longitudinal changes in PD.

Resting-state scans are often compromised by drowsiness and discomfort, even in healthy individuals [80, 81]. To address these prevalent confounds [40, 75, 82, 83], precision RSFC paradigms spread data collection across multiple sessions [39, 40, 53, 84]. Previous studies demonstrate, with sufficient data, functional networks are stable over time and exhibit minimal state-dependent variability, even across months or years [45, 46, 74, 85]. However, the equivalence of data across sessions had not previously been tested in PD. We find session-to-session similarity in the PD group is equivalent to that of healthy controls and previous work in healthy young adults [39]. Further, stability appeared to track with reliability (i.e., greater stability with more data per session) and the SNR of each structure (see **Supplemental Figure 2**). This is encouraging for the application of precision RSFC in PD to identify stable biomarkers. However, it raises questions about when network changes begin during the disease and whether they continue over time. Longitudinal studies indicate decreased RSFC over a few years in PD [86], which relates to decreased cognitive function [87]. Precision RSFC has not yet been used to study individual-level longitudinal changes in PD, marking this as an important next step.

### Individual differences in PD network topology

Group-level research on PD and RSFC has provided important insights into shared pathophysiology [15, 19] but is limited in its ability to explain clinical heterogeneity or inform precision care. Precision RSFC enables robust examination of interindividual variability [34] linked to behavior in healthy adults [48, 88, 89]. Although some work shows that RSFC networks based on group-level parcellations correlate with UPDRS-III scores in people with PD [90], there is a paucity of studies examining individualized networks. Here, we demonstrate it is possible to do so with precision methods. While group average network maps in PD aligned with those from healthy controls and young adults [39, 71, 91], PD participants differed from this group average and each other (**Figure 4**), consistent with past reports of individual variability [39, 46, 47, 92]. As a case example, a seed placed in the lateral motor cortex reveals robust somatomotor-face network connectivity in P001 and distinct somato-cognitive action network (SCAN) [72] connectivity in P026. These findings are again reflected in the distinct nature of cortical network parcellations in panel B. Our results highlight two major limitations in standard fMRI approaches in PD. First, methods assuming uniform network layouts across individuals risk blending signals from different networks, obscuring group-level interpretations. Second, standard approaches miss the opportunity to capture meaningful individual differences that may be linked to symptom profiles in PD.

### Future applications of Precision RSFC in PD

With the rise in personalized medicine, there is growing interest in understanding the mechanisms underlying heterogeneity in neurodegenerative diseases [93, 94]. Precision RSFC offers a powerful tool to link individual patterns of functional network organization to neuropathology and behavioral dysfunction. Its clinical potential has already been demonstrated in major depressive disorder, where network-level deviations (in this case, salience network expansion) predicted symptom severity [48]. Similar approaches could be applied in PD to better understand its broad spectrum of motor and non-motor symptoms.

Precision RSFC also holds promise in identifying personalized treatment targets. For instance, it could guide identification of cortical sites for transcranial magnetic stimulation or inform deep brain stimulation (DBS) planning by characterizing subcortical connectivity patterns. DBS outcomes vary greatly across targets [95] and individuals [96, 97] with evidence that variability in target connectivity contributes to differential efficacy [98–100]. For example, individual differences in globus pallidus connectivity, detectable through precision RSFC, correlates with outcome variability in DBS [50]. Ultimately, the RSFC patterns of therapeutic targets may serve as strong predictors of treatment efficacy, facilitating more precise and efficacious interventions in PD.

### Limitations

Several limitations should be acknowledged. This study included a relatively small sample (N = 26). However, the large amount of data per person in this precision RSFC approach produced highly reliable data, allowing for replication of results in each individual separately. Results were consistent across participants, and comparable in reliability, stability, and network organization with past precision RSFC work [39, 45], adding confidence in the findings. While larger samples may reveal group differences in motion, reliability, and stability between PD and HC, our results suggest such effects are likely modest. Additionally, participants were drawn from ongoing studies and may have shown greater tolerance for extended scanning, but the clinical similarity of our sample to those in previous PD RSFC studies [15] supports the generalizability of our results. In future studies recruiting MRI-naïve individuals, the use of a mock scanner or preliminary scan session to screen for contraindications (e.g., claustrophobia, excessive sleepiness, or high motion) may improve data quality. These strategies combined with the ability to collect data across multiple sessions, help mitigate many challenges that commonly impact data quality in PD fMRI research.

## Conclusion

Cumulatively, our findings suggest precision RSFC can be feasibly applied to studies of PD. Consistent with prior work in healthy populations, we replicate the finding that extended scan times substantially improve both the reliability and stability of RSFC metrics, outperforming those obtained from standard scanning protocols. PD participants achieved high test-retest reliability and day-to-day stability comparable to healthy controls. These results support the conclusion that network measures derived from precision RSFC in PD can meet the same rigorous standards established for healthy populations. More broadly, precision RSFC offers a powerful framework for identifying individualized network disruptions, with the potential to clarify the neural basis of clinical heterogeneity and to inform the development of personalized, network-targeted interventions.

## Supporting information

Supplemental_Fig1_motion

Supplemental_Fig2_tSNR

Supplemental_Fig3_reliability

Supplemental_Fig4_similarity

Supplemental_Table1_reliability

Supplemental_Table2_similarity

## Acknowledgments

We thank the participants for their time and effort with this project and the study coordinators for their diligent data collection. This work was supported by research funding from NIH (NS124738; NS075321; NS097437; NS097799), the Mallinckrodt Institute of Radiology and Neuroimaging Labs Research Center at WashU, the American Parkinson Disease Association, and the TOP TIER T32 program at WashU. This research was also supported in part by the computational resources and assistance from the staff at the Research Computing Center at Florida State University and the Illinois Campus Cluster. JC would also like to thank the Tallahassee Memorial Hospital Bryan W. Robinson Endowment.

## Author Roles

(1) Research Project: A. Conception, B. Organization. C. Execution (2) Statistical Analysis: A. Design, B. Execution, C. Review and Critique, (3) Manuscript Preparation:

A. Writing of the First Draft, B. Review and Critique J.C.: 1A, 1B, 1C; 2A, 2B; 3A

A.D.: 1C, 3A

S.G.: 1C; 3A

E.C.: 1C, 3A

A.E.: 1C; 3A

M.C.C.: 1A, 1B; 2A, 2C; 3B

C.G.: 1A, 1B; 2A, 2C; 3B.

