## Supplemental_Methods for "Towards Precision Functional Brain Network Mapping in Parkinson’s Disease"

### *fMRIprep*

The following boilerplate language describes the initial preprocessing steps completed via fMRIprep, edited to reflect the steps used in this manuscript: "Results included in this manuscript come from preprocessing performed using *fMRIPrep* 20.2.0 [1, 2], which is based on *Nipype* 1.5.1 [2, 3]. A total of 2 T1-weighted (T1w) images were found within the input BIDS dataset. All of them were corrected for intensity non-uniformity (INU) with N4BiasFieldCorrection [4], distributed with ANTs 2.3.3 [5]. The T1w-reference was then skull-stripped with a *Nipype* implementation of the antsBrainExtraction.sh workflow (from ANTs), using OASIS30ANTs as target template. Brain tissue segmentation of cerebrospinal fluid (CSF), white-matter (WM) and gray-matter (GM) was performed on the brain-extracted T1w using fast [6]. A T1w-reference map was computed after registration of 2 T1w images (after INU-correction) using mri_robust_template [7]. Brain surfaces were reconstructed using recon-all [8] and the brain mask estimated previously was refined with a custom variation of the method to reconcile ANTs-derived and FreeSurfer-derived segmentations of the cortical gray-matter of Mindboggle [9]. Volume-based spatial normalization to a standard space (MNI152NLin6Asym) was performed through nonlinear registration with antsRegistration (ANTs 2.3.3), using brain-extracted versions of both T1w reference and the T1w template. FSL's MNI ICBM 152 non-linear 6th Generation Asymmetric Average Brain Stereotaxic Registration Model [10] was used for spatial normalization.

For each of the BOLD runs found per subject (across all tasks and sessions), the following preprocessing was performed. First, a reference volume and its skull-stripped version were generated by aligning and averaging 1 single-band references (SBRefs). A B0-nonuniformity map (or *fieldmap*) was estimated based on a phase-difference map calculated with a dual-echo GRE (gradient-recall echo) sequence, processed with a custom workflow of *SDCFlows* inspired by the epidewarp.fslscript and further improvements in HCP Pipelines [11]. The fieldmap was then co-registered to the target EPI (echo-planar imaging) reference run and converted to a displacements field map (amenable to registration tools such as ANTs) with FSL's fugue and other *SDCflows* tools. Based on the estimated susceptibility distortion, a corrected EPI (echo-planar imaging) reference was calculated for a more accurate co-registration with the anatomical reference. The BOLD reference was then co-registered to the T1w reference using bbregister (FreeSurfer) which implements boundary-based registration [12]. Co-registration was configured with six degrees of freedom. Head-motion parameters with respect to the BOLD reference (transformation matrices, and six corresponding rotation and translation parameters) are estimated before any spatiotemporal filtering using mcflirt [13]. First, a reference volume and its skull-stripped version were generated using a custom methodology of *fMRIPrep*. The BOLD time-series (including slice-timing correction when applied) were resampled onto their original, native space by applying a single, composite transform to correct for head-motion and susceptibility distortions. These resampled BOLD time-series will be referred to as *preprocessed BOLD in original space*, or just *preprocessed BOLD*. The BOLD time-series were resampled into standard space, generating a *preprocessed BOLD run in MNI152NLin6Asym space*. A reference volume and its skull-stripped version were generated using a custom methodology of *fMRIPrep*.

All resamplings can be performed with *a single interpolation step* by composing all the pertinent transformations (i.e. head-motion transform matrices, susceptibility distortion correction when available, and co-registrations to anatomical and output spaces). Gridded (volumetric) resamplings were performed using antsApplyTransforms (ANTs), configured with Lanczos interpolation to minimize the smoothing effects of other kernels [14]. Non-gridded (surface) resamplings were performed using mri_vol2surf (FreeSurfer).

Many internal operations of *fMRIPrep* use *Nilearn* 0.6.2 [15], mostly within the functional processing workflow. For more details of the pipeline, see the section corresponding to workflows in fMRIPrep's documentation."

***FreeSurfer Edits***

T1-weighted scans were processed through FreeSurfer version 7.3 for cortical surface parcellation and subcortical volume segmentation. Within this pipeline, the input T1 images undergo normalization, skull stripping, white matter segmentation, cortical parcellation, and subcortical segmentation [16-18]. FreeSurfer output for each participant was manually visualized and manually edited to correct misclassifications (e.g., cerebellar and pial surface boundaries, extension of putamen into the claustrum) as necessary.
