## Supplemental_Fig1_motion for "Towards Precision Functional Brain Network Mapping in Parkinson’s Disease"

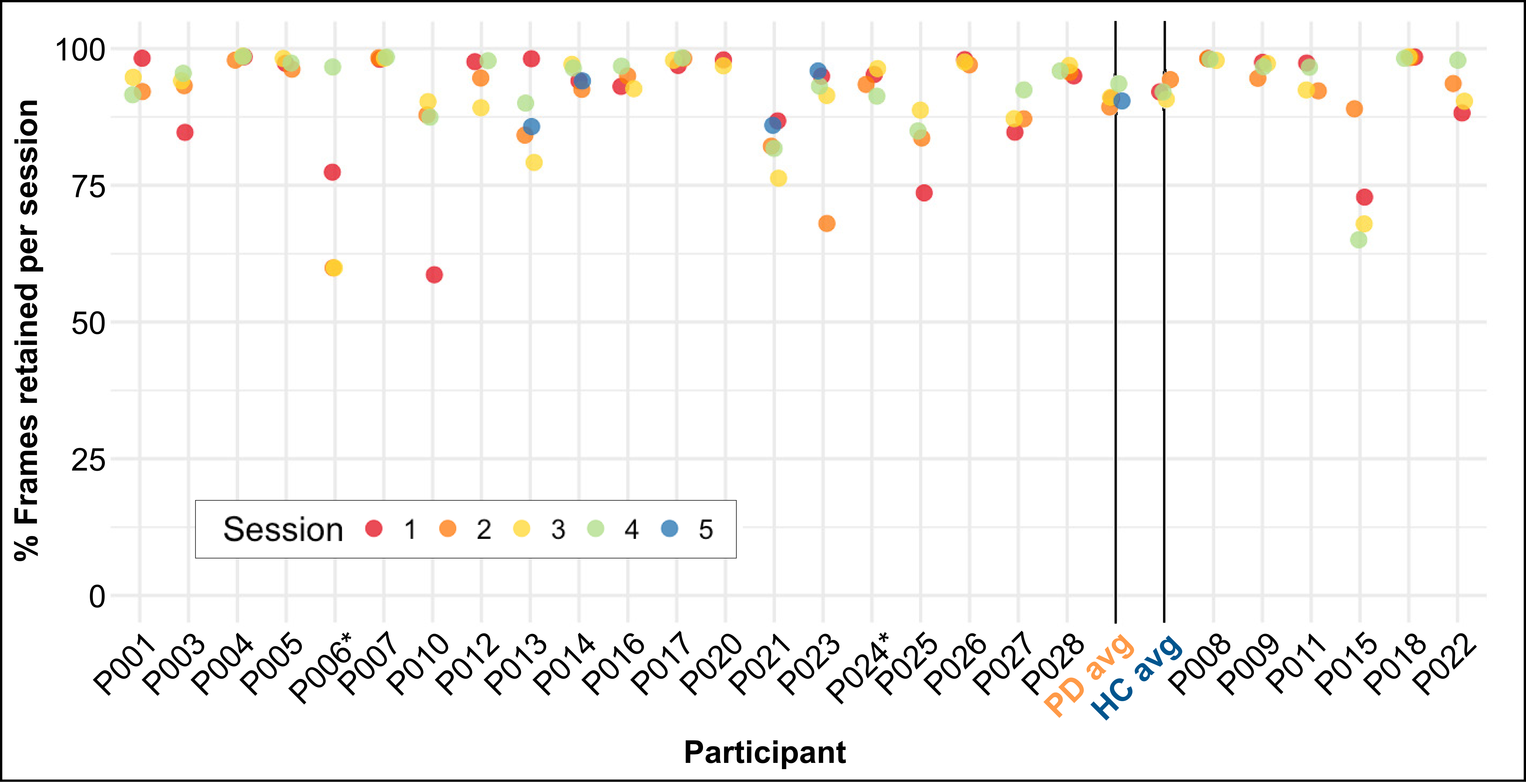


**Supplemental Figure 1: PD and HC participant data retention per session (different colors).** PD and HC participants retained and lost similar proportions of data due to motion in each session. For most participants, motion is relatively consistent across sessions. *Participant excluded from RSFC analyses due to a fieldmap acquisition error.
