## Supplemental_Fig2_tSNR for "Towards Precision Functional Brain Network Mapping in Parkinson’s Disease"

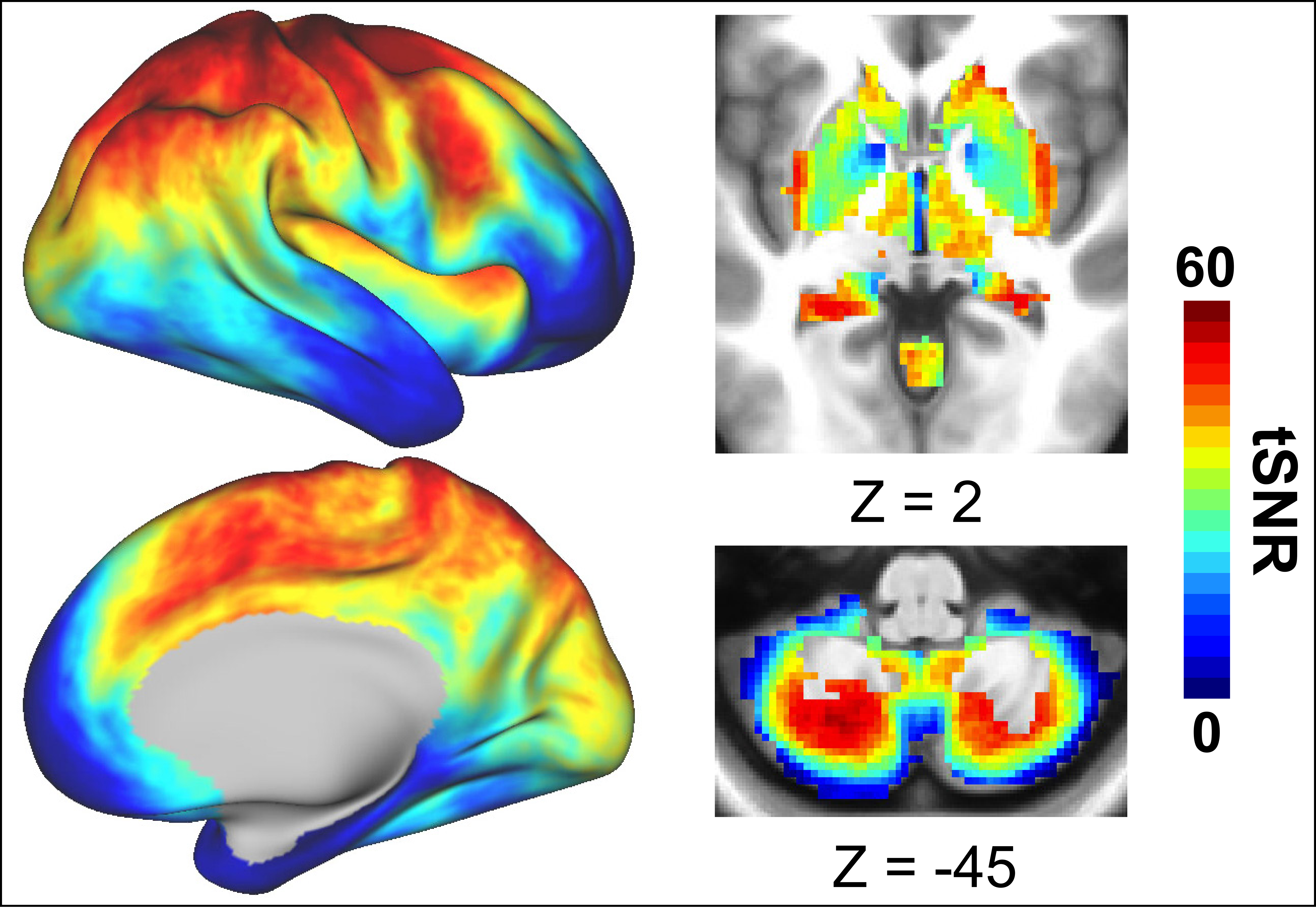


**Supplemental Figure 2: PD group average tSNR.** Following previous work [16, 17], tSNR diminishes as distance from the head coil increases. Consequently, the cortex registers the highest tSNR values, followed by the cerebellum, and the subcortex registers the lowest.
