## Supplemental_Fig3_reliability for "Towards Precision Functional Brain Network Mapping in Parkinson’s Disease"

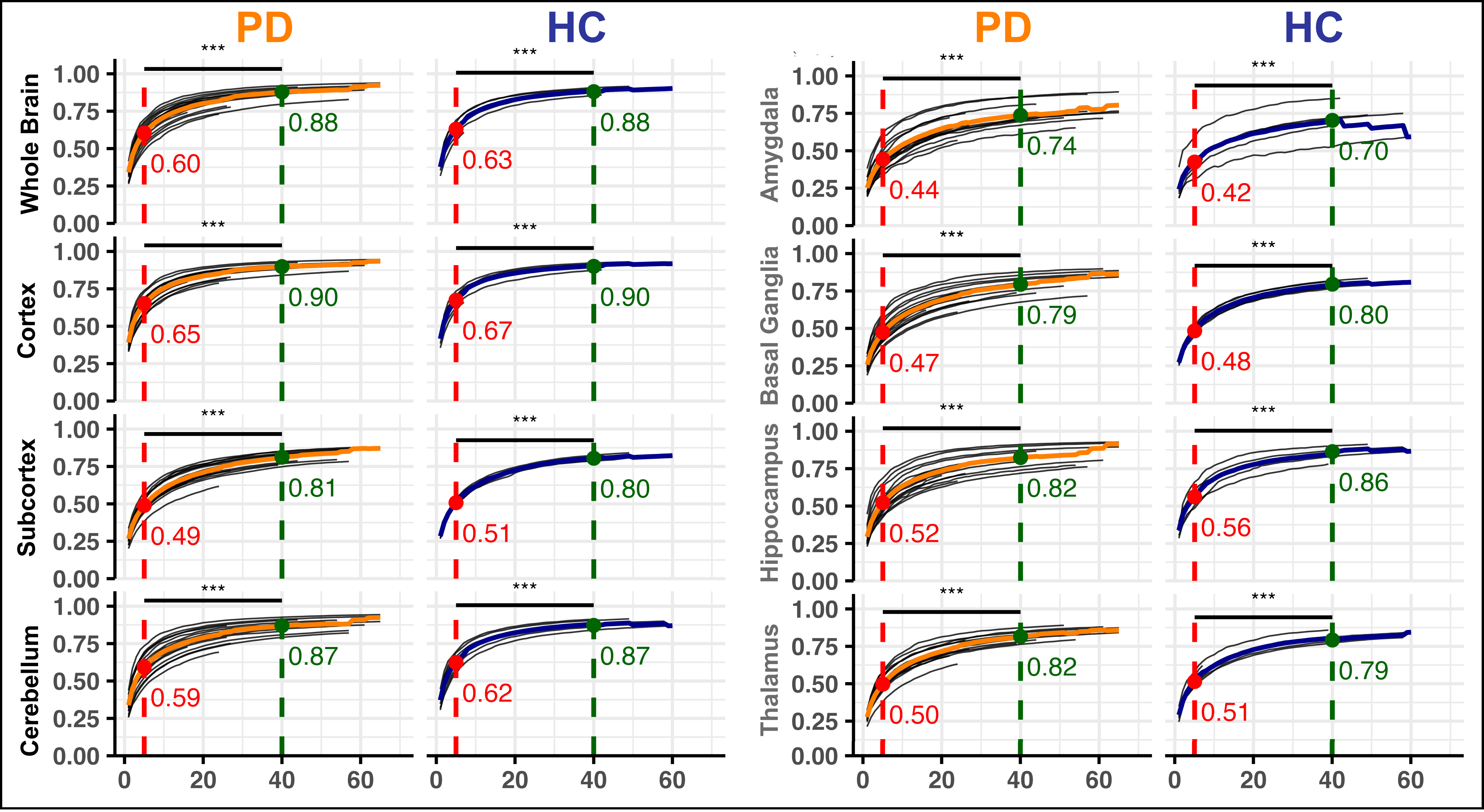


**Supplemental Figure 3: Precision fMRI Enhances Test-Retest Reliability of RSFC Differentially Across Brain Structures.** Test-retest reliability of RSFC improves significantly with increased scan time across all structures for both PD and HC groups ($p < .0001$ for all comparisons, see **Supp. Table 1**). As expected, subcortical regions, often affected by lower SNR, showed reduced reliability at 5 minutes but exhibited significant gains with longer scan durations, ultimately reaching good reliability levels. The subcortex includes the amygdala, basal ganglia, hippocampus, and thalamus.
