## Supplemental_Fig4_similarity for "Towards Precision Functional Brain Network Mapping in Parkinson’s Disease"

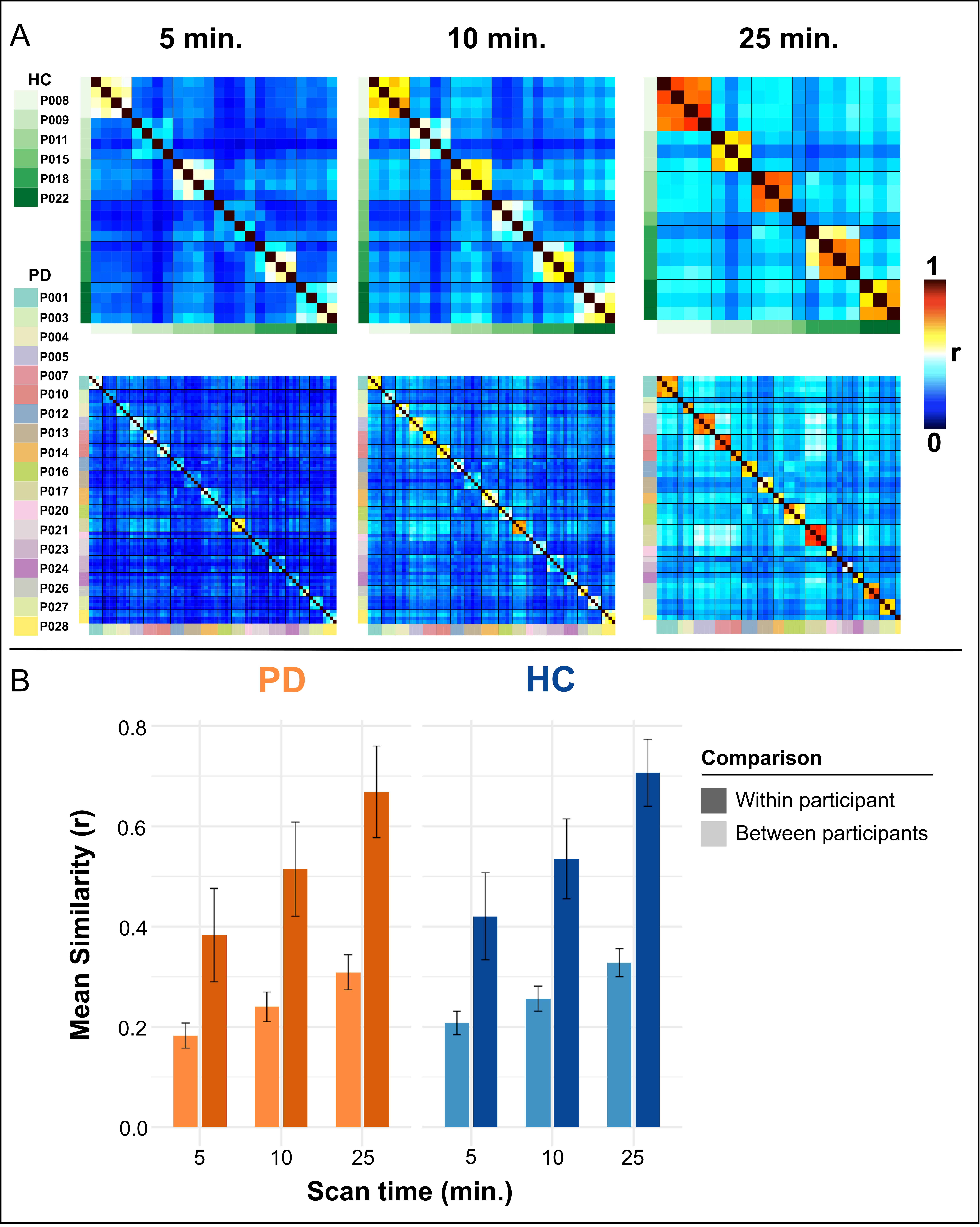


**Supplemental Figure 4: Session-to-Session RSFC Stability Improves with Increased Scan Time.** Mean similarity between RSFC matrices from separate scan days increased significantly with longer durations of low-motion data in both PD and HC participants. **A)** Each cell in the matrix reflects the average similarity in RSFC between connectivity profiles for a given amount of data (5, 10, and 25 min.) in each session. **B)** Paired t-tests showed robust gains in RSFC similarity with increasing scan time; Within-participant similarity (darker shades) was significantly higher than across-participant similarity (lighter shades) at all timepoints in both groups. For PD participants, significant increases in similarity were observed from 5 to 25 minutes ($t(14) = -24.99$, $p < 0.0001$) and from 10 to 25 minutes ($t(14) = -12.96$, $p < 0.0001$). For HC participants, similarity significantly increased from 5 to 25 minutes ($t(4) = -13.08$, $p = 0.0002$), and from 10 to 25 minutes ($t(4) = -10.02$, $p = 0.0006$). These results underscore the benefit of extended scan durations in improving test–retest reliability of functional connectivity patterns.
