## Supplementary figures and images for "Towards Precision Functional Brain Network Mapping in Parkinson’s Disease"

### Supplemental_Table1_reliability

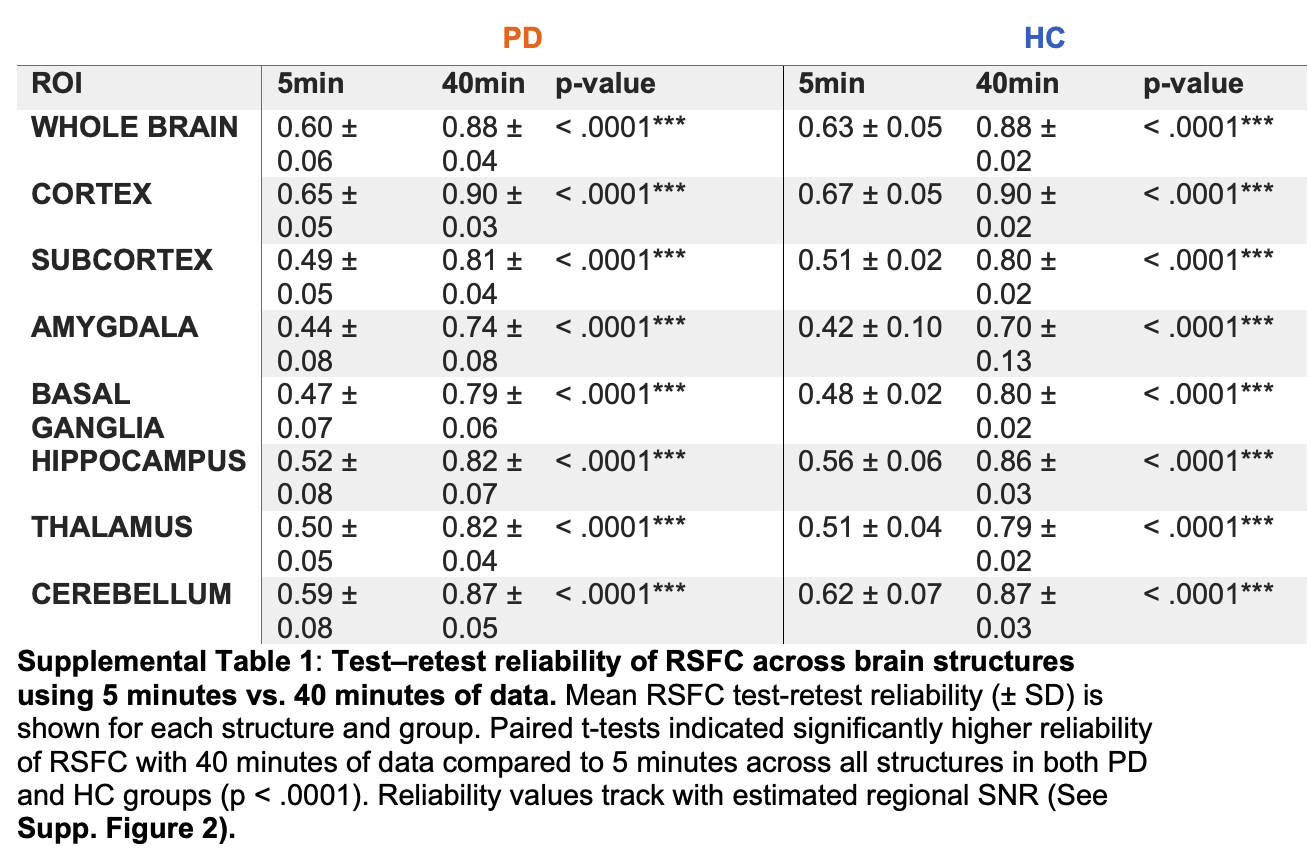

### Supplemental_Table2_similarity

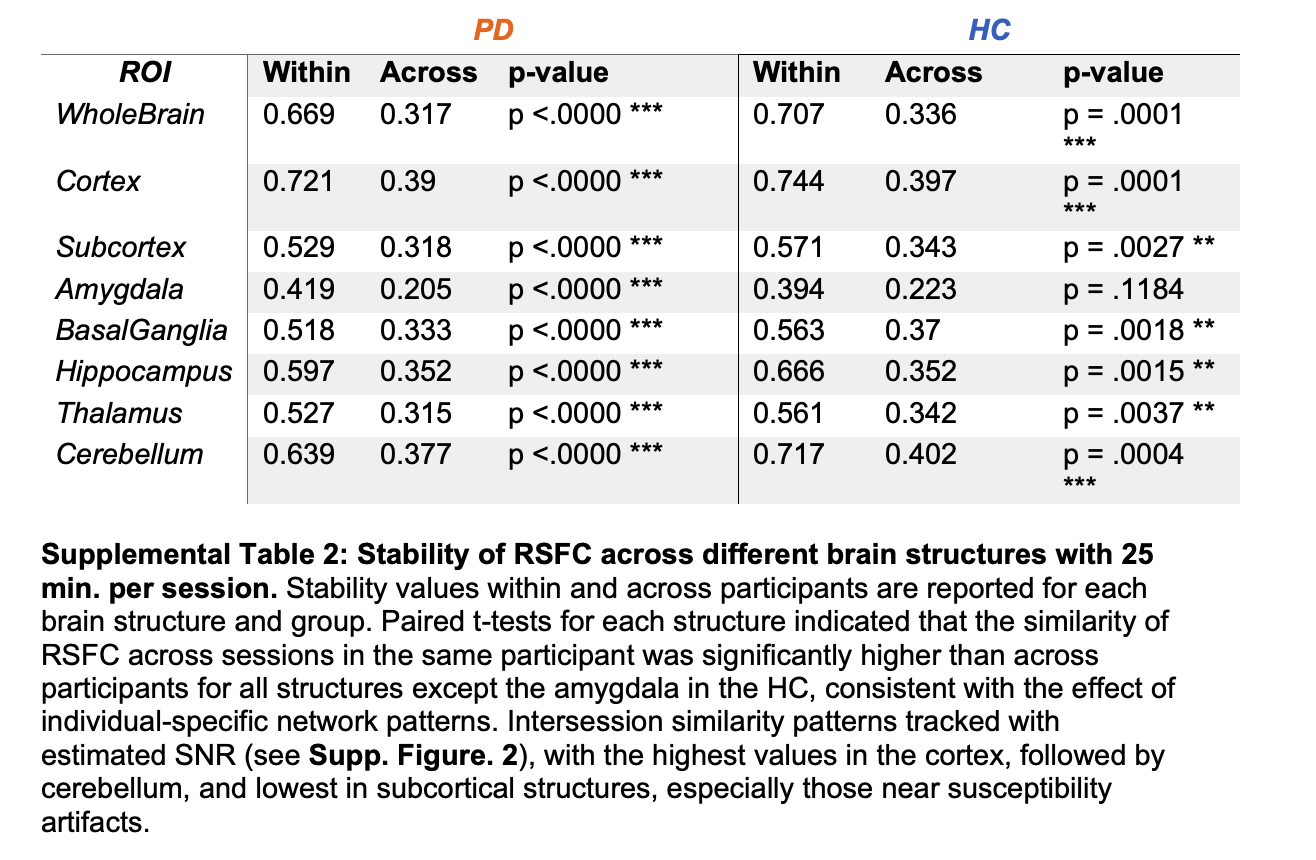
